# ReproPhylo: An Environment for Reproducible Phylogenomics

**DOI:** 10.1101/019349

**Authors:** Amir Szitenberg, Max John, Mark L. Blaxter, David H. Lunt

## Abstract

The reproducibility of experiments is key to the scientific process, and particularly necessary for accurate reporting of analyses in data-rich fields such as phylogenomics. We present ReproPhylo, a phylogenomic analysis environment developed to ensure experimental reproducibility, to facilitate the handling of large-scale data, and to assist methodological experimentation. Reproducibility, and instantaneous repeatability, is built in to the ReproPhylo system, and does not require user intervention or configuration because it stores the experimental workflow as a single, serialized Python object containing explicit provenance and environment information. This ‘single file’ approach ensures the persistence of provenance across iterations of the analysis, with changes automatically managed by the version control program Git. ReproPhylo produces an extensive human-readable report, and generates a comprehensive experimental archive file, both of which are suitable for submission with publications. The system facilitates thorough experimental exploration of both parameters and data. ReproPhylo is a platform independent CC0 python module, and is easily installed as a Docker image, with an Jupyter GUI, or as a slimmer version in a Galaxy distribution.

## Introduction

Experimental reproducibility has become a widely discussed issue in many areas of science [1,2]. Strict experimental reproducibility is not common in any area of the biological sciences and while the reasons for this may be varied they include the technical challenges in routine and robust implementation. Phylogenetic analyses are very widely used across the biological sciences [3], and, even in studies that are not primarily phylogenetic, the understanding of phylogenetic relationships is almost always required for a meaningful statistical inference [4–6]. Despite this importance, the reproducibility of phylogenetic experiments is low, and Magee et al. [7] estimated that 60% of published phylogenetic analyses are “lost to science” due the unavailability of the underlying data, an outcome also predicted in other areas of biology [8]. However, even the public archiving of all data does not ensure reproducibility, since complete knowledge of the analytical software, software versions, software parameters, dependencies and operating system versions can be very challenging to both discover and recreate from published manuscripts. The increasing quantity of DNA sequence data available, and the proliferation of analytic toolkits, makes phylogenetics carried out on a genomic scale (“phylogenomics”) both especially powerful, and especially problematic to reproduce. Reproducibility in phylogenomics requires tracking of data provenance of multiple loci from many taxa, and, frequently, deeply nested analyses that explore, sift and partition data to achieve the end goals of biological understanding.

Here we introduce ReproPhylo, a Python package designed to deliver reproducible phylogenomic analyses. ReproPhylo promotes reproducibility on two levels. First, it eases the complex phylogenomic pipeline design process by providing simple and concise scripting syntax for the execution of complex and forked phylogenetic workflows. Second, it automates reproducibility by employing well trusted containerization, versioning and provenance programs. In ReproPhylo, management of the experiment’s reproducibility and version control is carried out in a ‘frictionless’ manner in the background, without a need for user attention (although users have the option to access and tailor these aspects). With these two components of the analysis process considerably simplified, time and effort can be directed towards the core goals of understanding phylogenetic relationships by experimental parameter selection and data exploration, as the examples described here show (See Results section).

ReproPhylo is not the first package to provide phylogenetic workflow or pipeline tools [9–12]. A pipeline approach is a step forward from the point of view of reproducibility, as pipelines can serve as machine-readable records of analyses. Existing solutions [9–12] typically focus on the analysis itself, and do not attempt to provide complete reproducibility solutions. Several phylogenomic pipelines exist as web services [13–15], but server based analysis introduces additional complexities and reproducibility challenges. Osiris [16] achieves reproducibility through use of the Galaxy [17–19] reproducible bioinformatics environment. Within the Galaxy framework, Osiris offers tools and format converters for widely used phylogenetic analysis programs.

ReproPhylo explores an alternative, more generalised, approach to reproducibility. It unifies the different components of a flexible, convenient, platform-independent, user friendly and reproducible workflow, drawing on the many advantages of standard data formats and community standard Biopython [20] code classes. ReproPhylo is simply accessed within a Jupyter Notebook (formerly IPython Notebook) [21]. We have also designed several ReproPhylo Galaxy tools, which produce self-contained and fully reproducible outputs, even outside the Galaxy system, as a proof of concept.

## Design and Implementation

ReproPhylo interfaces with existing phylogenetic analysis tools via standard data structures, such as 

~~~
SeqRecord
~~~

 or 

~~~
MultipleSeqAlignment
~~~

 Biopython objects. In addition, it imports and exports data as text files in all standard formats supported by Biopython [20], and does not itself implement any novel data formats.

ReproPhylo can be run using Jupyter Notebook [21] as a command line interface or a GUI. We provide a range of notebooks for different types of analysis with the ReproPhylo distribution, including one for the Leptidoptera case analysis presented below. These notebooks are examples of ‘literate programming’ [22] in that they combine instructions, documentation, and code. The user may modify these Notebook pipelines either trivially (e.g. just changing the input data and executing), or more substantially (by altering the nature or sequence of analyses via Python code). Our testing with undergraduates, postgraduates, and academics without coding experience indicates that Jupyter Notebook is an effective GUI for scientists lacking a background in programming.

## The ReproPhylo pipeline

ReproPhylo aids processes through the complete arc of a phylogenomics study: dataset collation, data analysis and visualisation/ exploration. Table 1 lists the data classes in ReproPhylo and their associated methods and functions. The ReproPhylo module uses a set of Python packages to control the pipeline and report results and quality statistics. The workflow is carried out by Biopython [20] and ETE2 [23], the latter of which also powers tree annotation. The primary output data file format is PhyloXML, although other formats can be produced. Graphics other than phylogenetic trees, such as alignment statistics and sequence statistics box-plots, are produced using Matplotlib [25].

Dataset collation in ReproPhylo has three components: harvesting, selection and filtering. An example of data harvest would be importing all GenBank records for a specific taxonomic group from a Genbank format text file, and adding unpublished sequences from a fasta or ab1 format sequence file. Exonerate [26] can be deployed within ReproPhylo to harvest loci of interest from genome or transcript data via specialized functions. Data selection exploits ReproPhylo’s loci report to automatically include or exclude specific genes and coding sequences present in an input Genbank file. Data filtering automatically excludes or includes sequences, or loci, based on user specifications - length, GC content, sequence number or taxonomic coverage - informed by ReproPhylo’s sequence and alignment summary statistics reports.

The analysis workflow in ReproPhylo includes sequence alignment, alignment trimming, and tree reconstruction. These steps can be forked to explore alternative analytic approaches while tracking data provenance in each branch and step. We have included commonly used analytic tools for each step, and additional algorithms can be suggested or included using Python code, as detailed in the manual (http://goo.gl/yW6J1J). The first release of ReproPhylo can utilise the sequence aligners MAFFT [27], MUSCLE [28,29] and Pal2Nal [30]. Trimming of alignments to remove poorly aligned ‘gappy’ regions can improve analyses [31], and is carried out based on explicit trimming criteria using TrimAl [32]. Tree reconstruction programs accessible through ReproPhylo include RAxML PTHREADS SSE3 [33] and PhyloBayes [34].

ReproPhylo facilitates phylogenetic output visualisation and exploration. Tree annotation, and creation of publication quality figures, is powered by ETE2 [23] and informed by metadata from the data harvest step provided to it by ReproPhylo. Bayestraits [35,36] is included for comparative phylogenetic analyses, and is invoked by a function which accepts a ReproPhylo Project object as the source of both the tree and trait information. Pairwise tree distances between trees in the 

~~~
Project
~~~

 can be computed and visualized (see Results section).

**Table 1:**
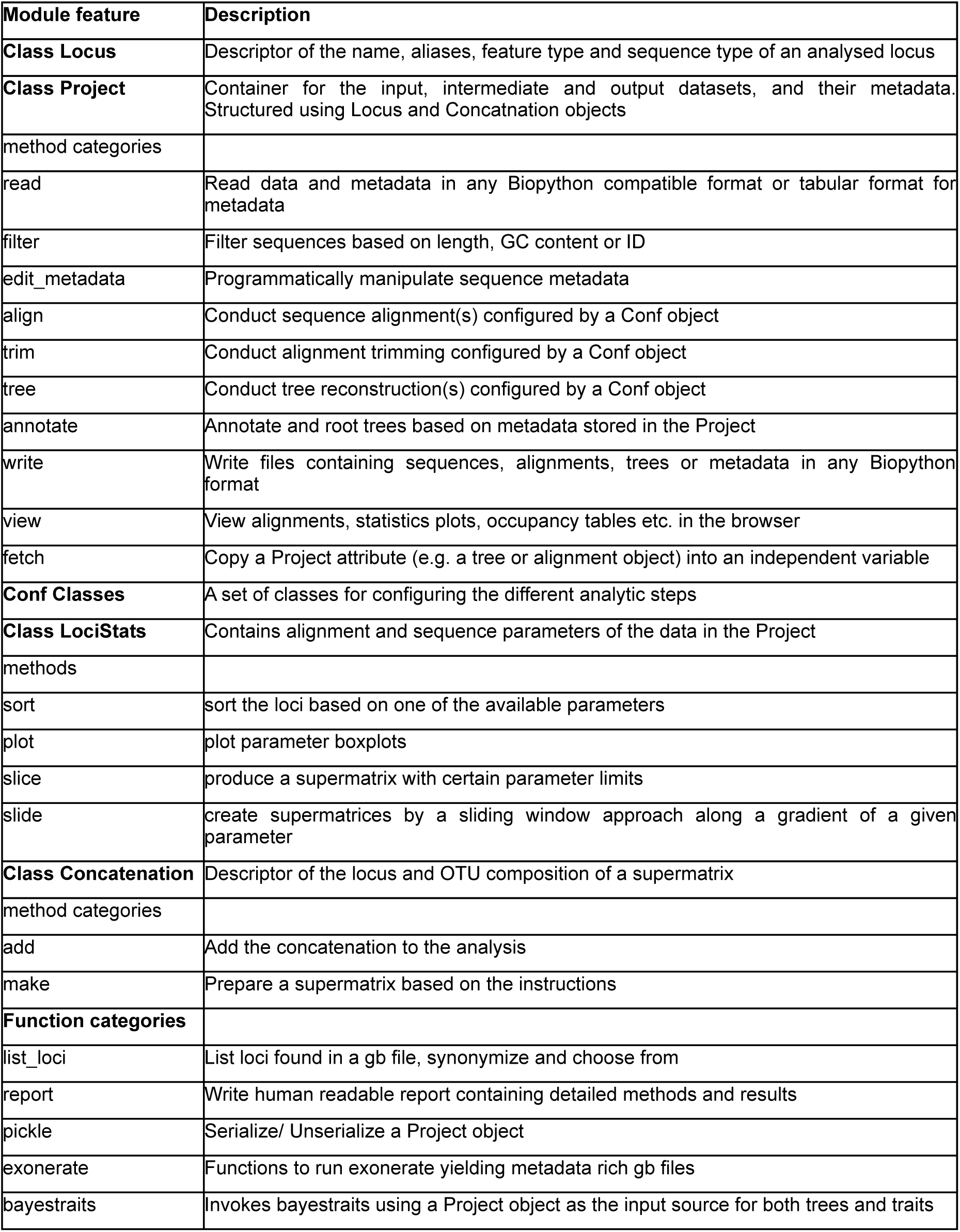
Summary of the the Python module structure

## Data provenance and reproducibility

Data provenance, the recording of the input and transformation of information used to generate a result, is a key issue in reproducibility. To maintain phylogenomic data provenance, ReproPhylo keeps the full workflow in a single instance of the Project ReproPhylo class (Figure 1A). This object contains all the analytical steps and their outputs, together with machine and human readable unique process IDs that describe the provenance of each data object for both the program and the user. In addition, the Project instance contains the metadata associated with each sequence of each locus, with a unique ID, which allows it to associate the metadata with its sequence or tree leaf in any of the existing data objects (the 

~~~
SeqRecord, MultipleSeqAlignment
~~~

 and 

~~~
Tree
~~~

 objects). Analysis is invoked by Project class methods, which modify the data (e.g. align the sequences), place the resulting data object (e.g. 

~~~
MultipleSeqAlignment
~~~

) in the appropriate Project attribute (e.g. Project.alignments) under an unique ID (Figure 1B), update the binary file storing the Project, and commit it to the Git repository. In each analytical step metadata can be retrieved using unique sequence identifiers, and alternative analytic approaches (forks) can be stored within a single Project through their unique process IDs.

**Fig. 1.**
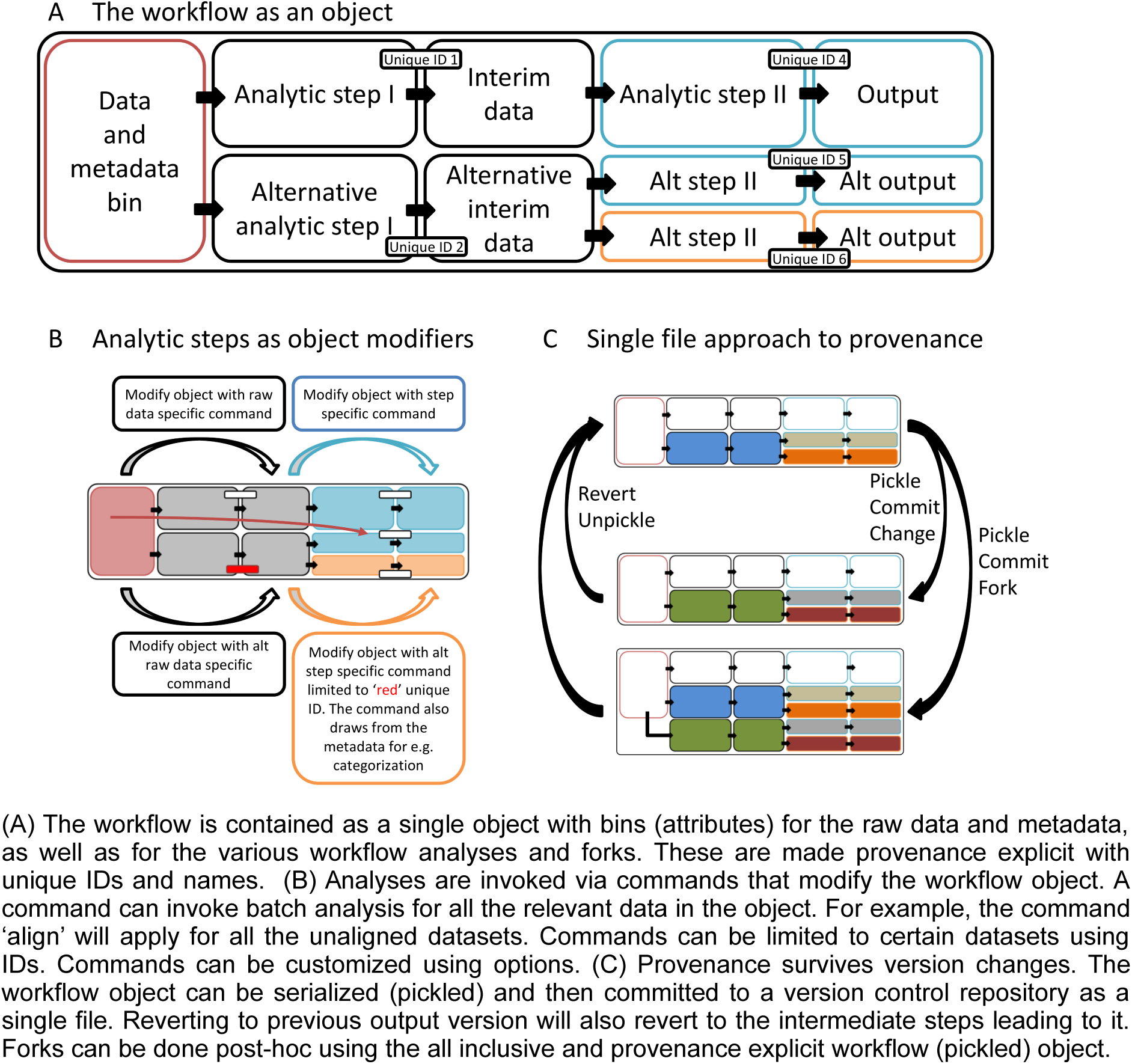
The phylogenetic workflow as a single python object. (A) The workflow is contained as a single object with bins (attributes) for the raw data and metadata, as well as for the various workflow analyses and forks. These are made provenance explicit with unique IDs and names. (B) Analyses are invoked via commands that modify the workflow object. A command can invoke batch analysis for all the relevant data in the object. For example, the command ‘align’ will apply for all the unaligned datasets. Commands can be limited to certain datasets using IDs. Commands can be customized using options. (C) Provenance survives version changes. The workflow object can be serialized (pickled) and then committed to a version control repository as a single file. Reverting to previous output version will also revert to the intermediate steps leading to it. Forks can be done post-hoc using the all inclusive and provenance explicit workflow (pickled) object.

Since the complete workflow is represented as a single Python object, provenance can be maintained across different versions of the analysis (Figure 1C). ReproPhylo serializes (“pickles”) the Project object and maintains it as a binary file that allows the user to pause and resume the analysis seamlessly. ReproPhylo uses the version control program Git (git-scm.com) to record a version of the binary Project file each time it is modified, and thus allows forwards and backwards toggling of file versions. When an older version is restored, the full chain of intermediate results and the records detailing their production are restored throughout the workflow and across forks. ReproPhylo’s version control and reproducibility are implemented passively in the background and are frictionless for the user, requiring neither specialist knowledge or action to produce a reproducible phylogenomics experiment.

To facilitate publication of the reproducible experiment, ReproPhylo produces a compressed experiment directory (.zip format) suitable for upload to a data depository such as FigShare (http://figshare.com/) or DataDryad (http://datadryad.org/). This file contains trees and sequence alignments (in standard PhyloXML format), all analysis scripts, tree figure files, and a complete, human-readable report. The report includes a methods section ready for inclusion in a manuscript, which contains program versions, accession numbers, references etc., to which the digital object identifier of the full experimental record can be added. The compressed experiment directory also contains the binary file in which the serialized Project object is stored. This object contains all the data, metadata, method descriptions and results, and includes explicit provenance information. It can be used to revive the entire analysis, either in a Docker container, in a local ReproPhylo installation or independently of ReproPhylo, and instantly repeat it or extend it. Another product of ReproPhylo is a Git repository, which can be published on websites such as Github (http://github.com/) and Figshare (http://figshare.com/).

## The ReproPhylo pipeline

ReproPhylo is open source, using strictly open source dependencies, and is under active development within a publicly accessible Github repository (https://github.com/HullUnibioinformatics/ReproPhylo). Documentation is provided as a version tracked publicly-editable Google Docs manual at http://goo.gl/yW6J1J, allowing corrections and expansions by the user community. A frozen version of the module (Version 1), utilizing Jupyter Notebook as interface, is available as a self contained environment in a Docker image (http://goo.gl/JcHMGN). Bioinformatics pipelines may often be challenging to install but the use of a Docker image (http://www.docker.com) for distribution eliminates such difficulties, and facilitates installation on any system. The Docker image is accompanied by a shell script that will install and deploy the ReproPhylo image as a Docker container, with a local web browser based GUI. Using Docker as a work environment also facilitates reproducibility and reusability, as all relevant files can be committed to the image, generating a single Docker image file containing the computer environment, specific program copies, and data components of the finished analysis. Such containerisation approaches, which deliver both reproducible and easily reusable experiments, are powerful development and delivery tools. As a proof of concept, ReproPhylo is also provided as a Galaxy distribution (http://goo.gl/udsS3Q) containing ReproPhylo Galaxy tools. This version utilises the Galaxy framework, while retaining completely reproducible results even outside the Galaxy GUI.

## Demonstration

Several examples of use of the ReproPhylo phylogenomic analytical pipeline are provided as Jupyter notebooks in the distribution files. We focus here on parameter space exploration using ReproPhylo to demonstrate the advantages to phylogenomic analysis delivered by a fully scripted, reproducible environment. In this use case we demonstrate exploration of the effect of the median residue conservation (gene variability level) in each locus on a resulting species topology, using an existing, multigene dataset from species in Lepidoptera [37]. Loci with different levels of conservation may hold phylogenetic signal of events that occurred in different times in the past, or may be too conserved, or may be too rapidly evolving and saturated with homoplasies, to provide any signal at all [38]. We utilise Shannon Entropy (SE) [39] as a conservation scoring method [40]. The script generating this analysis is available as Methods S1 (http://dx.doi.org/10.6084/m9.figshare.1409427). The original Jupyter Notebook, together with the input and output files and figures, has been archived in FigShare (doi:10.6084/m9.figshare.1409423), and has also been included as one of the tutorials in the current distribution of ReproPhylo (see documentation at http://goo.gl/aZeRXf). A report with supplementary results generated by ReproPhylo is provided as Results S1. Instructions on accessing the Project file in order to reproduce this demonstration are provided in the manual.

We obtained a nucleotide sequence alignment of 465 loci from 26 Lepidoptera species [37]. Using a built-in function (Methods S1, section 2.6.1), SE values [39], ignoring gap characters, were calculated for each residue in each locus. An entropy distribution plot (Figure 2A, centre) illustrates the differences in SE among the loci. This plot is typical of alignment statistics and representations produced by the ReproPhylo 

~~~
LociStats
~~~

 class (see Section 2.6.3 of Methods S1 for code generating this plot). Six supermatrices were extracted, each from a sliding window of 200 loci, starting with the highest entropy loci and ending with the lowest entropy loci, and shifting the window by 50 loci between subsets (Figure 2A). Lastly, following the original analysis, all 26 species were included in all of the supermatrices, which contained no missing data (Results S1, section 2.7).

**Fig. 2.**
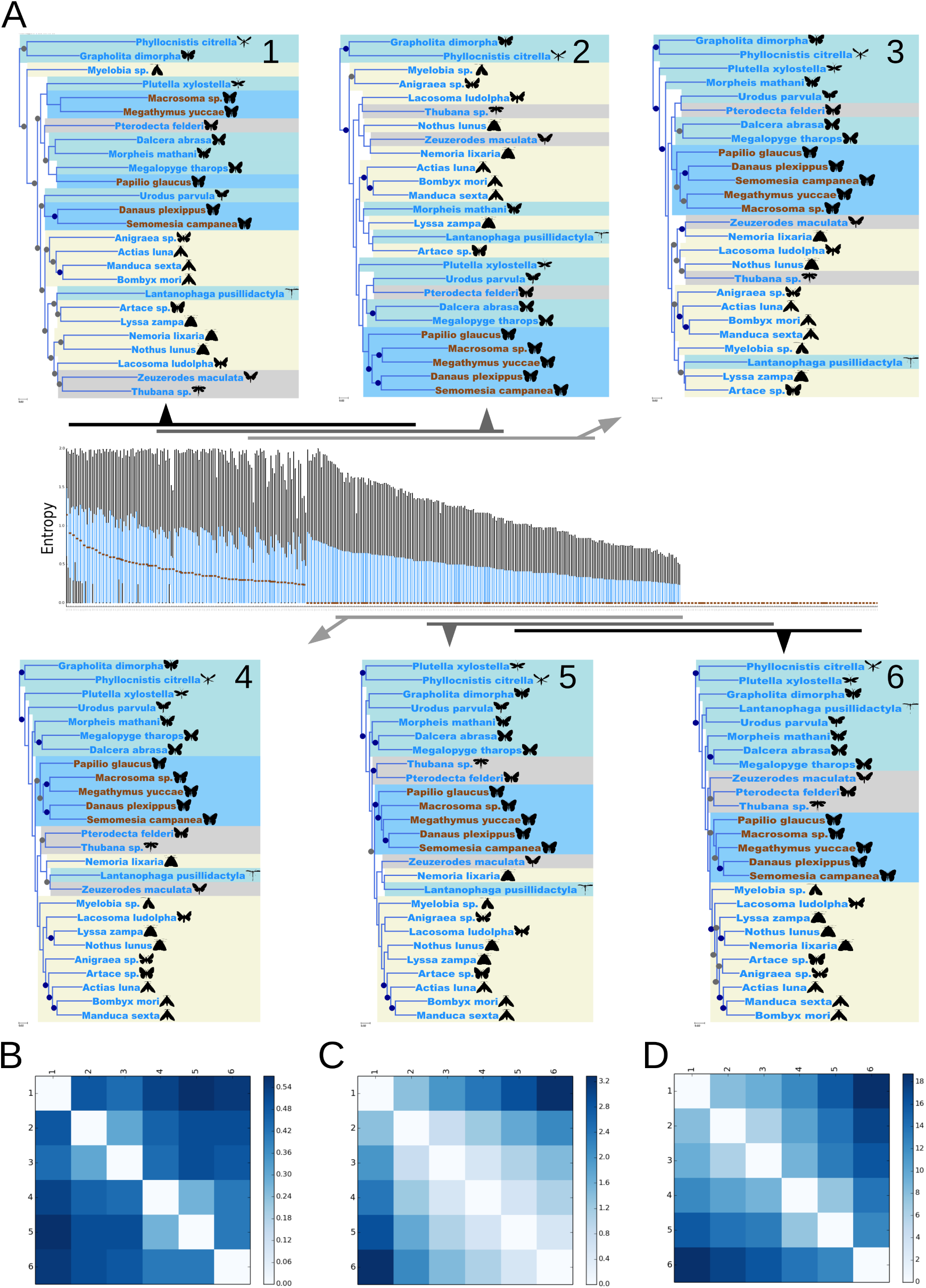
Exploratory phylogenomic analysis of a Lepidoptera dataset. (A) A nucleotide dataset from 26 species from Kawahara and Breinholt [37] was reanalyzed. Loci were sorted by their median, 75 percentile and 25 percentile entropy values (centre panel). For each locus, a box plot was generated. The medians are denoted by brown dots. The boxes (blue) represent the 25-75 percentiles. Whiskers (black) represent values that are found within a range outside the box, 1.5 times as long as the box (which is null, when the box itself has a null range) Trees (insets A 1 - 6) were reconstructed from 200-locus windows with 50 locus overlap between neighbouring windows. The windows are represented by black and gray horizontal bars, each with an arrow pointing to the tree generated from it. In trees 1 - 6, dark blue highlights denote Rhopalocera (butterfly) taxa, and light blue, gray and yellow highlights denote clades I, III and IV respectively (sensu Kawahara and Breinholt [37]). Bullets on nodes represent Bootstrap percentages (BP). Blue bullets represent maximal support. Other support values above 80% are denoted by gray bullets. (B-D) Three pairwise tree divergence metrics were calculated and presented as heatmaps, with the most divergent tree pairs denoted by dark blue and identical tree pairs by a white box. While the scales are not comparable among the metrics, the relative differences are. The metrics are (B) the Symmetric Distance of Robinson-Foulds [41], (C) the Branch Distance [42] and (D) evolutionary rate corrected Branch Distance [42].

While ReproPhylo provides a sequence alignment step, the alignments were retained as originally published. They were trimmed to exclude any positions composed entirely of gaps, using the 

~~~
TrimalConf
~~~

 class, which invokes TrimAl [32] (Methods S1, section 2.5). Trees were reconstructed with RAxML 8 [33], invoked using the 

~~~
RaxmlConf
~~~

 class, which automatically passes a partition file. RAxML was configured to execute a single ML tree search with 100 rapid bootstrap replicates, using the GTRGAMMA model (see Methods S1, section 2.8, and Results S1). 

~~~
RaxmlConf
~~~

 allows the user to set models specifically for each partition, or set of partitions, but we used a single model here to standardise across locus partition sets. The trees produced are presented as Figure 2A, insets 1-6. The script that generated these trees is in section 2.10 of Methods S1.

To formally compare phylogenetic trees, ReproPhylo can calculate three metrics (Methods S1, section 2.11) and plot them as heat maps, where the most divergent tree pairs are indicated with dark blue blocks and identical trees by white ones (Figure 2 B-D). The first, the Symmetric Distance of Robinson-Foulds [41] (Figure 2B), accounts for topological differences alone, while the second, the Branch Distance [42, 43] (Figure 2C), also incorporates branch-length information. The third, a modified Branch Distance [42], (Figure 2D) attempts to standardize evolutionary rates across trees.

## Reproducibility statement

The entire project workflow was saved as a pickle file (Results S1), the Git repository generated by ReproPhylo (doi:10.6084/m9.figshare.1409423) and a publishable archive file (Results S1). The pickled workflow can most productively be used within the ReproPhylo environment, where it is possible to add data and repeat the analysis or extend the analysis without the need to repeat any previous step. Importantly, the data within the pickled workflow is accessible using Biopython, even in the absence of ReproPhylo. The archive file represents a more traditional approach to reproducibility, as it includes alignment and tree text files, the tree figures (Figure 2B) and a human readable report containing complete methods and results information.

## Results

We explored the partitioned Lepidoptera data for support for the clade Rhopalocera (butterflies) in loci with different SE values. Butterfly taxa are indicated in Figure 2B with dark blue highlight. High entropy loci provided insufficient signal to support the monophyly of Rhopalocera, or to resolve relationships among its lineages (Figure 2A, tree 1). Loci with intermediate entropy values contain strong signal for the phylogenetic relationships within the Rhopalocera (Figure 2A, trees 2-4), and loci with low entropy values provide support for monophyly, but are insufficient to resolve the internal relationships, as evidenced by the topology and node support values (Figure 2A trees 5-6). For three other clades identified by Kawahara and Breinholt [37] (their clades I, III and IV; Figure 2A insets, light blue, yellow and gray highlights respectively), monophyly is only recovered in the lowest entropy dataset (Figure 2A tree 6). The entropy calculations were shown to be unbiased by the GC content, sequence length and gap-score distributions of the loci, in Figure S1 (generated by the code in section 2.4.6, Methods S1).

Trees that were created by different branches of the workflow were formally compared using tree distance metrics. Corrected Branch Distance (Figure 2D) better reflects topological differences between trees than Branch Distance (Figure 2C), but still accounts for the signal supporting bifurcations, while Symmetric Distance does not (Figure 2B). Symmetric Distance comparison revealed profound topological differences among the different partitions. When branch lengths were considered (Figure 2C, Branch Distance), these differences were more graded, with trees from neighbouring sliding windows more similar than other tree pairs, a result that was expected given the influence of entropy on branch lengths. When evolutionary rates are standardized (Figure 2D, corrected Branch Distance), profound topological differences among the different partitions are again revealed.

Overall, we found that the topology was not robust to variation in residue entropy, and thus affirm that any study should carefully document and justify locus selection. The key novelty in the ReproPhylo environment is the ease and flexibility with which a complex phylogenetic investigation such as this can be set up, and be instantaneously repeatable and reproducible without compromising the user’s control over parameter choice and configuration. ReproPhylo facilitates informed parameter choices and data filtering based on clearly documented and reproducible experimentation. Additional use cases are included with the package and they demonstrate the usage of additional components of the module and their interaction with Git and Docker.

## Conclusions

ReproPhylo is an integrated environment for performing fully reproducible, platform independent, phylogenomics analyses that is highly accessible for scientists even without a strong computational background. ReproPhylo, by dealing with input and output formatting of data and results, can improve the accessibility and integration of existing computational tools. ReproPhylo is intended to be a community tool, and we hope its future development will be guided by input from users, either by pull requests or issue reporting and suggestions in the Github repository.

Phylogenetic analyses focussing on a single locus are becoming rarer as the power of modern genomics makes the de novo generation of large-scale data for multiple species feasible, especially with targeted sequencing approaches [44]. The rapid growth of public databases provides a resource that can be mined for new sets of loci across wide taxonomic spans, offering a second source of very large phylogenomic datasets. To exploit these new data, and at the same time deliver fully reproducible science that can deliver to a truly incremental synthesis of evolution of life on earth, toolkits such as ReproPhylo that are large-data-ready, and natively reproducible will be essential.

## Supplementary material

http://dx.doi.org/10.6084/m9.figshare.1409426

**Figure S1**. **Loci statistics boxplots for data derived from** [37].

For each locus, the plots illustrate the distributions of (from top to bottom) per-position entropy, per-position gap score [32], per position conservation score [32], sequence length and GC content. http://dx.doi.org/10.6084/m9.figshare.1409424

**Methods S1**. A static HTML representation of the code that was used to create all the analyses in this study.

**nbviewer:** http://goo.gl/KzFAvj

http://dx.doi.org/10.6084/m9.figshare.1409427

**Results S1**. A results archive produced by ReproPhylo, containing the serialized Project, input and output files, scripts and an HTML report.

http://dx.doi.org/10.6084/m9.figshare.1409488

## Acknowledgements

We thank Dr. Africa Gómez, Dr. Christoph Hahn, Stephen Moss, Daniel Jefferies, and Claudia Scavariello for useful comments on the program and the manuscript. AS, DL and MB are supported in part by NERC award NE/J011355/1. The GenePool has core support from the NERC (award R8/H10/56) and MRC (G0900740).

The silhouettes in Figure 2 are distributed here under the Creative Commons Attribution 3.0 Unported and are credited as follows:

Geometroidea, Bombycoidea: Gareth Monger

Cossoidea: Didier Descouens (vectorized by T. Michael Keesey) Gelechioidea: Caroline Harding, MAF (vectorized by T. Michael Keesey)

## References

1. McNutt M. Journals unite for reproducibility. Science. 2014;346: 679.

2. Begley CG, Ioannidis JPA. Reproducibility in science improving the standard for basic and preclinical research. Circ Res. 2015;116: 116–126.

3. Eales JM, Pinney JW, Stevens RD, Robertson DL. Methodology capture: discriminating between the “best” and the rest of community practice. BMC Bioinformatics. 2008;9: 359.

4. Penny D. The comparative method in evolutionary biology. J Classification. 1992;9: 169–172.

5. Whitney KD, Baack EJ, Hamrick JL, Godt MJW, Barringer BC, Bennett MD, et al. A role for nonadaptive processes in plant genome size evolution? Evolution. 2010;64: 2097–2109.

6. Ågren JA, Wang W, Koenig D, Neuffer B, Weigel D, Wright SI. Mating system shifts and transposable element evolution in the plant genus Capsella. BMC Genomics. 2014;15: 602.

7. Magee AF, May MR, Moore BR. The dawn of open access to phylogenetic data. PLoS ONE. 2014;9: e110268.

8. Vines TH, Albert AYK, Andrew RL, Débarre F, Bock DG, Franklin MT, et al. The availability of research data declines rapidly with article age. Curr Biol. 2014;24: 94–97.

9. Huerta-Cepas J, Bork P, Gabaldon T. ETE-NPR: A portable application for Nested Phylogenetic Reconstruction and workflow design. Submitted.

10. Pearse WD, Purvis A. phyloGenerator: an automated phylogeny generation tool for ecologists. Methods Ecol Evol. 2013;4: 692–698.

11. Grant JR, Katz LA. Building a phylogenomic pipeline for the eukaryotic tree of life - addressing deep phylogenies with genome-scale data. PLoS Curr. 2014;6. doi:10.1371/currents.tol.c24b6054aebf3602748ac042ccc8f2e9

12. Dunn CW, Howison M, Zapata F. Agalma: an automated phylogenomics workflow. BMC Bioinformatics. 2013;14: 330.

13. Sánchez R, Serra F, Tárraga J, Medina I, Carbonell J, Pulido L, et al. Phylemon 2.0: a suite of web-tools for molecular evolution, phylogenetics, phylogenomics and hypotheses testing. Nucleic Acids Res. 2011;39: W470–4.

14. Dereeper A, Guignon V, Blanc G, Audic S, Buffet S, Chevenet F, et al. Phylogeny.fr: robust phylogenetic analysis for the non-specialist. Nucleic Acids Res. 2008;36: W465–9.

15. Miller MA, Wayne P, Terri S. Creating the CIPRES Science Gateway for inference of large phylogenetic trees. 2010 Gateway Computing Environments Workshop (GCE). 2010. doi:10.1109/gce.2010.5676129

16. Oakley TH, Alexandrou MA, Ngo R, Pankey MS, Churchill CKC, Chen W, et al. Osiris: accessible and reproducible phylogenetic and phylogenomic analyses within the Galaxy workflow management system. BMC Bioinformatics. 2014;15: 230.

17. Giardine B, Riemer C, Hardison RC, Burhans R, Elnitski L, Shah P, et al. Galaxy: A platform for interactive large-scale genome analysis. Genome Res. 2005;15: 1451–1455.

18. Blankenberg D, Kuster GV, Coraor N, Ananda G, Lazarus R, Mangan M, et al. Galaxy: A web-based genome analysis tool for experimentalists. Current Protocols in Molecular Biology. John Wiley & Sons, Inc.; 2001.

19. Goecks J, Nekrutenko A, Taylor J. Galaxy: a comprehensive approach for supporting accessible, reproducible, and transparent computational research in the life sciences. Genome Biol. 2010;11: R86.

20. Cock PJA, Antao T, Chang JT, Chapman BA, Cox CJ, Dalke A, et al. Biopython: freely available Python tools for computational molecular biology and bioinformatics. Bioinformatics. 2009;25: 1422–1423.

21. Pérez F, Granger BE. IPython: a system for interactive scientific computing. Comput Sci Eng. 2007;9: 21–29.

22. Knuth DE. Literate programming. Comput J. 1984;27: 97–111.

23. Huerta-Cepas J, Dopazo J, Gabaldón T. ETE: a python environment for tree exploration. BMC Bioinformatics. 2010;11: 24.

24. Sukumaran J, Holder MT. DendroPy: a Python library for phylogenetic computing. Bioinformatics. 2010;26: 1569–1571.

25. Hunter JD. Matplotlib: A 2D graphics environment. Comput Sci Eng. 2007;9: 90–95.

26. Slater GSC, Birney E. Automated generation of heuristics for biological sequence comparison. BMC Bioinformatics. 2005;6: 31.

27. Katoh K, Standley DM. MAFFT multiple sequence alignment software version 7: improvements in performance and usability. Mol Biol Evol. 2013;30: 772–780.

28. Edgar RC. MUSCLE: multiple sequence alignment with high accuracy and high throughput. Nucleic Acids Res. 2004;32: 1792–1797.

29. Edgar RC. MUSCLE: a multiple sequence alignment method with reduced time and space complexity. BMC Bioinformatics. 2004;5: 1–19.

30. Suyama M, Torrents D, Bork P. PAL2NAL: robust conversion of protein sequence alignments into the corresponding codon alignments. Nucleic Acids Res. 2006;34: W609–W612.

31. Talavera G, Castresana J. Improvement of phylogenies after removing divergent and ambiguously aligned blocks from protein sequence alignments. Syst Biol. 2007;56: 564–577.

32. Capella-Gutiérrez S, Silla-Martínez JM, Gabaldón T. trimAl: a tool for automated alignment trimming in large-scale phylogenetic analyses. Bioinformatics. 2009;25: 1972–1973.

33. Stamatakis A. RAxML Version 8: A tool for phylogenetic analysis and post-analysis of large phylogenies. Bioinformatics. 2014; btu033.

34. Lartillot N, Lepage T, Blanquart S. PhyloBayes 3: a Bayesian software package for phylogenetic reconstruction and molecular dating. Bioinformatics. 2009;25: 2286–2288.

35. Pagel M. Detecting correlated evolution on phylogenies: a general method for the comparative analysis of discrete characters. Proc R Soc B. 1994;255: 37–45.

36. Pagel M, Meade A, Barker D. Bayesian estimation of ancestral character states on phylogenies. Syst Biol. 2004;53: 673–684.

37. Kawahara AY, Breinholt JW. Phylogenomics provides strong evidence for relationships of butterflies and moths. Proc R Soc B. 2014;281: 20140970.

38. Higgs PG. RNA secondary structure: physical and computational aspects. Q Rev Biophys. 2000;33: 199–253.

39. Shannon CE. A Mathematical Theory of Communication. SIGMOBILE Mob Comput Commun Rev. 2001;5: 3–55.

40. Valdar WSJ. Scoring residue conservation. Proteins. 2002;48: 227–241.

41. Robinson DF, Foulds LR. Comparison of phylogenetic trees. Math Biosci. 1981;53: 131–147.

42. Kuhner MK, Felsenstein J. A simulation comparison of phylogeny algorithms under equal and unequal evolutionary rates. Mol Biol Evol. 1994;11: 459–468.

43. Sukumaran J, Holder MT. DendroPy: a Python library for phylogenetic computing. Bioinformatics. 2010;26: 1569–1571.

44. Lemmon AR, Emme SA, Lemmon EM. Anchored hybrid enrichment for massively high-throughput phylogenomics. Syst Biol. 2012;61: 727–744.

